# Pharmacologic analysis of non-synonymous coding 5-HT2A SNPs reveals alterations psychedelic drug potencies and efficacies

**DOI:** 10.1101/2021.12.09.472000

**Authors:** Gavin P. Schmitz, Manish K. Jain, Samuel T. Slocum, Bryan L. Roth

## Abstract

Serotonin (5-Hydroxytryptamine; 5-HT) 2A receptor (5-HT_2A_R) signaling is essential for the actions of classical psychedelic drugs. In this study, we examined whether random sequence variations in the gene (single nucleotide polymorphisms, SNPs) encoding the 5-HT_2A_R affect the signaling of four commonly used psychedelic drugs. We examined the *in vitro* pharmacology of seven non-synonymous SNPs, which give rise to S12N, T25N, D48N, I197V, A230T, A447V, and H452Y variant 5-HT_2A_ serotonin receptors. We found that these non-synonymous SNPs exert statistically significant, although modest, effects on the efficacy and potency of four therapeutically relevant hallucinogens. Significantly, the *in vitro* pharmacological effects of the SNPs drug actions at 5-HT_2A_R are drug specific.

**Table of Contents/Abstract Graphic:** 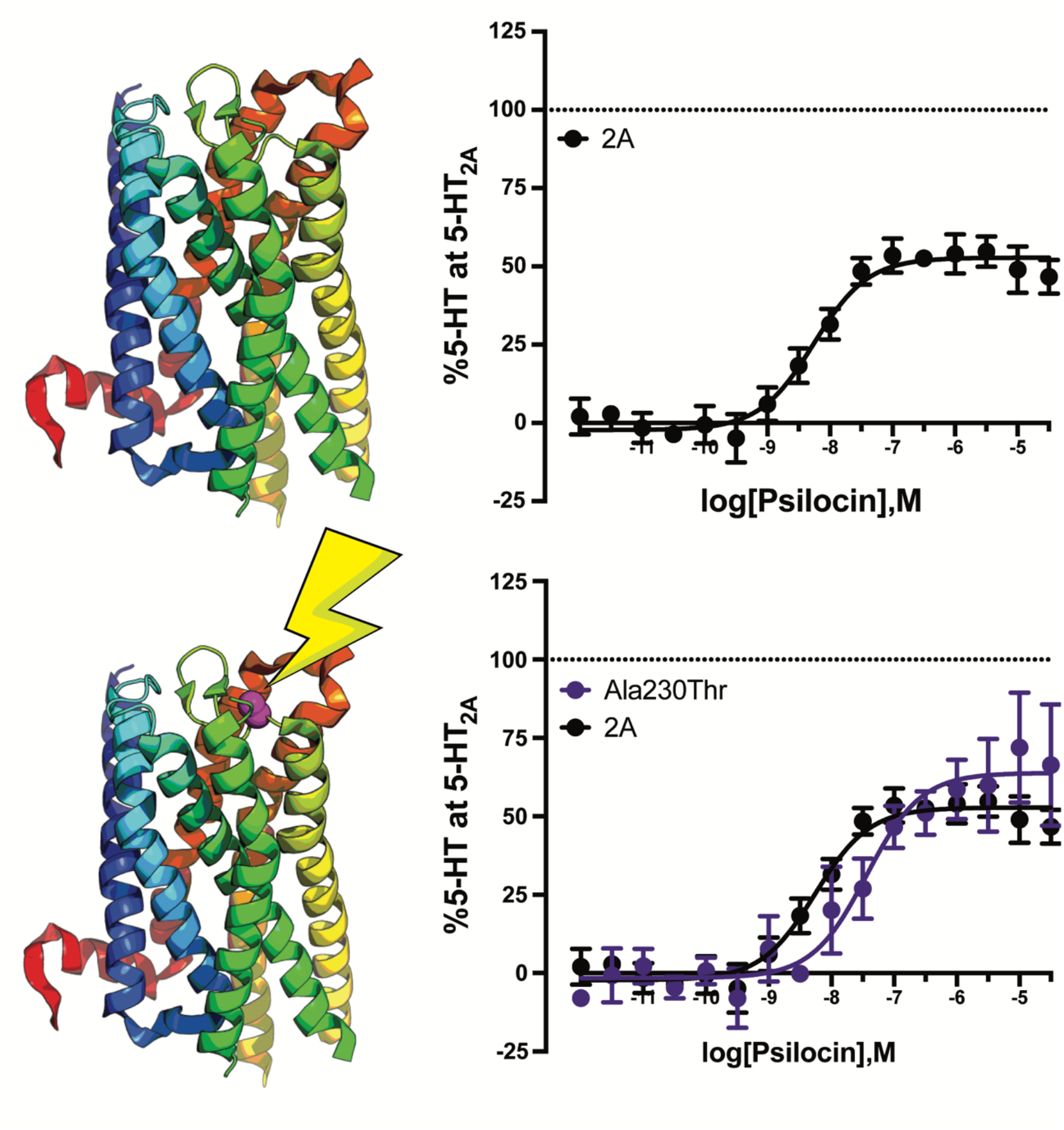

## 1. Introduction

Psychedelic drugs have been defined as compounds that induce lysergic acid diethylamide (LSD)-like effects on cognition, emotion, and perception via activation of 5-HT_2A_ serotonin (5-hydroxytryptamine; 5-HT) receptors^1, 2^. Recently, there has been renewed interest in psychedelic compounds as potential therapeutics for many neuropsychiatric disorders^3^. For example, psilocybin, the prodrug to the active compound psilocin found in *Psilocybe cubensis* mushrooms, has recently been granted breakthrough drug status by the Food and Drug Administration due to its potential as a treatment for treatment-resistant depression and anxiety^4, 5, 6^. These effects of psilocybin in these Phase II clinical trials are both rapid and apparently enduring^4, 5, 7^. Similarly, LSD has demonstrated efficacy in treating cluster headaches^8^ and alleviating anxiety in terminally ill patients^9^. Finally, anecdotal reports and recent meta-analyses have suggested the potential therapeutic utility for mescaline and 5-methoxy-*N,N*-dimethyltryptamine (5-MeO-DMT) in treating anxiety, depression, and other affective disorders^10–12, 13, 14^.

Psychedelics are generally classified by chemical structure (e.g. tryptamines; ergolines and phenylisopropylamines) and each psychedelic drug has a robust pharmacology, with activities at many serotonin and other biogenic amine receptors^15–19^. 5-HT_2A_ receptor agonism has been shown to be crucial in mediating the psychoactive effects of psychedelics in both animal^20–22^ and human studies^23–26^. 5-HT_2A_ receptors are members of the G protein-coupled receptors (GPCR) superfamily and canonically signal through the Gα_q_ family of heterotrimeric G proteins activating phospholipase Cβ and many other down-stream effector systems ^27, 28, 2^. 5-HT_2A_ receptors also recruit β-arrestin (βArr) proteins *in vitro*^29^ and the 5-HT_2A_R is apparently complexed with βArr1 and βArr2 in cortical neurons *in vivo*^30^. While many GPCR agonists activate both G protein- and βArr-mediated signaling pathways, certain ligands are known to preferentially activate one signaling pathway via a ligand-dependent phenomenon known as functional selectivity or biased signaling^31^. It has been proposed that ligands with biased profiles may be useful for activating desired pathways (i.e. therapeutically efficacious) rather than undesired pathways (i.e. unwanted side effect producing) downstream of a given GPCR^32^. Indeed, LSD is shown to preferentially signal through βArr2 at the 5-HT_2A_ receptor^18, 33^ and numerous LSD-elicited responses are either significantly attenuated or completely absent in βArr2-KO mice^34^.

Early clinical studies on psilocin have shown a wide variability in drug response^35^ with a significant portion of respondents showing no statistically significant treatment effect. The reasons for such inter-individual variability in hallucinogenic drug response are unknown. At the molecular level, random sequence variations in genes (single nucleotide polymorphisms, SNPs) could explain inter-individual differences in drug response and such activities are highly relevant as hallucinogens become more prevalent in clinical practice. At least seven non-synonymous SNPs are located within the coding region of the human 5-HT_2A_ receptor gene (HTR2A, 13q14-21) (Fig. 1). These seven single-nucleotide polymorphisms (S12N, T25N, D48N, I197V, A230T, A447V, and H452Y) have potential actions on both receptor structure and function^36^. As shown in Figure 1, the S12N, T25N, and D48N SNPs map to the predicted N-terminal tail of the receptor; the I197V SNP resides in the fourth transmembrane helix; the A230T SNP resides in the putative second extracellular loop (ECL2); the two remaining non-synonymous SNPs (A447V and H452Y) are located within the putative C-terminal tail. These SNPs represent seven of the most common variants observed in human populations with allele frequencies varying from 0.003% (T25N) to 7.9% (H452Y) (https://gpcrdb.org/)^37^.

**Figure 1:**
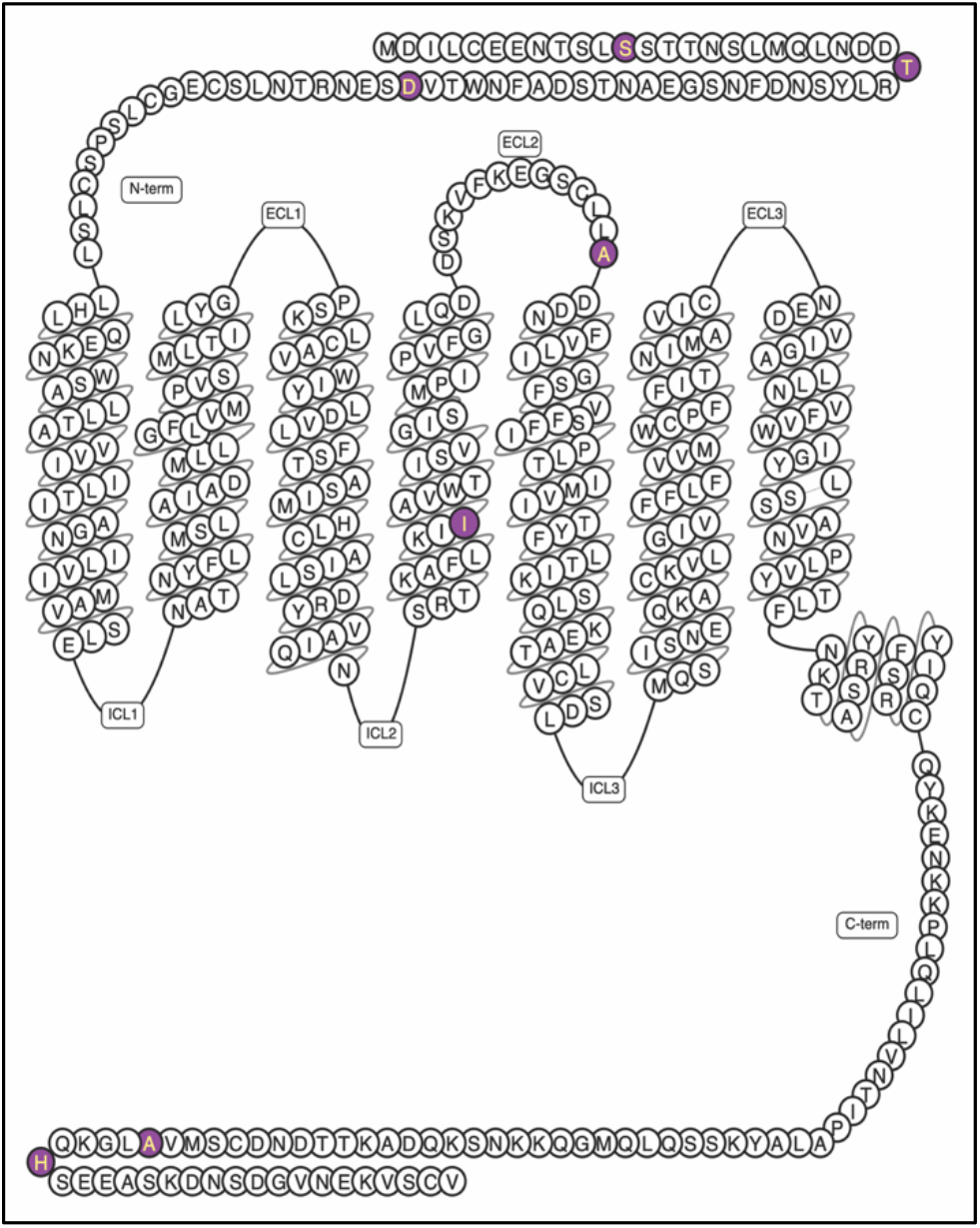
Snake plot representation of the 5-HT_2A_ receptor showing the location of the seven SNPs: S12N, T25N, D48N, I197V, A230T, A447V, H452Y.

Here, we comprehensively examined the potency and signaling bias for four therapeutically promising psychedelics at several 5-HT_2A_ non-synonymous SNPs. We discovered that the effects of individual SNPs were drug selective and that no single SNP had uniform effects on all psychedelics tested. These results imply that future clinical studies into the therapeutic utility of psychedelics might consider particular drug-SNPs combinations – paying particular attention to those SNPs that cause significant differences in the *in vitro* pharmacologies of test agents.

## 2. Materials and Methods

### 2.1 Cell culture

HEK293T cells were obtained from ATCC (Manassas, VA). Cells were maintained, passaged, and transfected in Dulbelcco’s Modified Eagle’s Medium containing 10% fetal bovine serum (FBS), 100 Units/mL penicillin, and 100μg/mL streptomycin (Gibco-ThermoFisher, Waltham, MA) in a humidified atmosphere at 37°C and 5% CO2. After transfection, cells were plated in DMEM containing 1% dialyzed FBS, 100 Units/mL penicillin, and 100μg/mL streptomycin for BRET assays.

### 2.2 Mutagenesis

The human 5-HT_2A_ cDNA cloned into the *Not*I site pIRES-neo (Clonetech) was obtained through the resources of the National Institute of Mental Health Psychoactive Drug Screening Program (http:kidb.pdsp.unc.edu). The following mutagenesis primers were obtained from Eton Bioscience:

S12N forward: 5’ TCACTGAACAGCACAACTAACTCTCTCATG 3’

S12N reverse: 5’ TGTGCTGTTCAGTGAGGTATTCTCCTC 3’

T25N forward: 5’ GATGATAACCGACTTTACTCCAACGACTTCAAC 3’

T25N reverse: 5’ AAGTCGGTTATCATCATTGAGCTGCATGAG 3’

D48N forward: 5’-ACAGTAAACAGTGAAAACCFFACCAATCTGTCC

D48N reverse: 5’-TTCACTGTTTACTGTCCAATTGAAGGCGTC

I197V forward: 5’ AAAATCGTCGCGGTCTGGACCATTTCA 3’

I197V reverse: 5’ GACCGCGACGATTTTCAGAAAGGCTTT 3’

A230T forward: 5’ CTTCTGACTGATGACAATTTCGTACTTATAGGAAGC 3’

A230T reverse: 5’ GTCATCAGTCAGAAGGCAACTGCCCTC 3’

A447V forward: 5’ ATGGTGGTCCTGGGCAAACAGCACAGTGAAGAAGCC 3’

A447V reverse: 5’ GCCCAGGACCACCATACTGCAGTCGTTGTC 3’

H452Y forward: 5’ AAACAGTACAGTGAAGAAGCCTCCAAAGAC 3’

H452Y reverse: 5’ TTCACTGTACTGTTTGCCCAGGGCCAC 3’

SNPs were introduced into the cloned human 5-HT_2A_ receptor template using the QuikChange site directed, PCR-based mutagenesis kit exactly as described by the manufacturer (Stratagene, La Jolla, CA, USA). Mutagenized clones were isolated and subjected to Sanger DNA sequencing to verify the entire coding sequence for the presence of the desired mutation and the absence of any PCR-induced sequence errors.

### 2.3 BRET2 assays

Cells were plated either in six-well dishes at a density of 700,000–800,000 cells/well, or 10-cm dishes at 7–8 million cells/dish. Cells were transfected 2–4 hours later, using a 1:1:1:1 DNA ratio of receptor:Gα-RLuc8:Gβ:Gγ-GFP2 (100 ng/construct for six-well dishes, 1000 ng/construct for 10-cm dishes). Transit 2020 (Mirus Biosciences, Madison, WI) was used to complex the DNA at a ratio of 3 μL Transit/μg DNA, in OptiMEM (Gibco-ThermoFisher, Waltham, MA) at a concentration of 10 ng DNA/μL OptiMEM. The next day, cells were harvested from the plate and plated in poly-D-lysine-coated white, clear bottom 96-well assay plates (Greiner Bio-One, Monroe, NC) at a density of 30,000–50,000 cells/well.

One day after plating in 96-well assay plates, white backings (Perkin Elmer, Waltham, MA) were applied to the plate bottoms, and growth medium was carefully aspirated and replaced immediately with 50 μL of assay buffer (1x HBSS + 20 mM HEPES, pH 7.4) containing freshly prepared 50 μM coelenterazine 400a (Nanolight Technologies, Pinetop, AZ). After a five-minute equilibration period, cells were treated with 50 μL of drug (2X) for an additional 5 minutes. Plates were then read in a Pherastar FSX microplate reader (BMG Labtech Inc., Cary, NC) with 395 nm (RLuc8-coelenterazine 400a) and 510 nm (GFP2) emission filters, at 1 second/well integration times. BRET2 ratios were computed as the ratio of the GFP2 emission to RLuc8 emission. Results are from at least three independent experiments, each performed in duplicate. Data were normalized to 5-HT stimulation and analyzed using nonlinear regression “log(agonist) vs. response” in Prism 9 (Graphpad Software Inc., San Diego, CA).

### 2.4 Bioluminescence Resonance Energy Transfer (BRET) Arrestin Assay

To measure 5-HT_2A_R-mediated β-arrestin1 recruitment, HEK293T cells were co-transfected in a 1:1:5 ratio with human 5-HT_2A_R containing C-terminal *Renilla* luciferase (*R*Luc8), GRK2, and Venus-tagged N-terminal β-arrestin1. To measure 5-HT_2A_R-mediated β-arrestin2 recruitment, HEK293T cells were co-transfected in a 1:1:5 ratio with human 5-HT_2A_R containing C-terminal *Renilla* luciferase (*R*Luc8), GRK2, and Venus-tagged N-terminal β-arrestin2. For both experiments, after at least 24 hours, transfected cells were plated in poly-lysine coated 96-well white clear bottom cell culture plates in plating medium (DMEM + 1% dialyzed FBS) at a density of 40,000–50,000 cells in 200 µL per well and incubated overnight. The next day, medium was decanted, and cells were washed with 50 µL of drug buffer (1X HBSS, 20 mM HEPES, 0.1% BSA, 0.01% ascorbic acid, pH 7.4), then 50 µL of drug buffer with *R*Luc substrate coelenterazine h (Promega, 5 µM final concentration) was added per well. After 5 minutes, 50 µL of drug (2X) was added per well and incubated for 5 minutes. Plates were read for both luminescence at 485 nm and fluorescent eYFP emission at 530 nm for 1 second per well using a Pherastar FSX microplate reader (BMG Labtech Inc., Cary, NC). The ratio of eYFP/*R*Luc was calculated per well and the net BRET ratio was calculated by subtracting the eYFP/*R*Luc per well from the eYFP/*R*Luc ratio in wells without Venus-β-Arrestin present. Results are from at least three independent experiments, each performed in duplicate. The net BRET ratio was plotted as a function of drug concentration using Prism 9 (Graphpad Software Inc., San Diego, CA). Data were normalized to 5-HT stimulation and analyzed using nonlinear regression “log(agonist) vs. response” in Prism 9.

### 2.5 Receptor Expression Measurements

HEK293T cells were plated in 10cm dishes and transfected with the wild-type 5-HT_2A_ R, S12N, T25N, D48N, I197V, A230T, A447V, or H452Y variant of 5-HT_2A_ receptor. After 36h of transfection, cell medium was replaced with medium containing 1% dialyzed FBS. After 48 h, medium was aspirated and cells were washed twice with cold phosphate-buffered saline (PBS). Cells were scraped in 10 mL ice cold PBS and centrifuged at 3000 rpm at 4C for 15 minutes and the supernatant was discarded. The cell pellet was resuspended in ice cold lysis buffer (50 mM Tris HCl, pH 7.4) and incubated on ice for 20 min. After incubation, cells were triturated gently for hypotonic lysis. The suspension was centrifuged at 21000g for 20 minutes at 4C to obtain a crude membrane pellet. The supernatant was discarded, and the membrane pellets were stored at −80°C for further use.

Saturation binding assays were performed to determine equilibrium dissociation constant (Kd) and Bmax. Membrane pellets were dissolved in lysis buffer and the amount of protein was estimated using Bradford assays. Binding assays were performed in standard binding buffer (50 mM Tris, 10 mM MgCl2, 0.1 mM EDTA, 0.1% BSA, 0.01% ascorbic acid, pH 7.4) using [^3^H] ketanserin (PerkinElmer, Waltham, MA; specific activity 22.8 Ci/mmol) as the radioligand. Saturation binding assays were carried out in 96-well plates in a final volume of 125 μL per well. In brief, 25 μL of radioligand was added to a 96-well plate followed by addition of 25 μL binding buffer (for total binding) or 25 μL reference compound (Clozapine) at a final concentration of 10 μM to measure nonspecific binding. After, 75 μL of fresh membrane protein (15 μg/well) was added and plates were incubated in the dark at room temperature for 90 minutes. The reaction was stopped by vacuum filtration onto cold 0.3% polyethyleneimine (PEI)-soaked 96-well filter mats using a 96-well Packard Filtermate harvester, followed by three washes with cold wash buffer (50 mM Tris, pH 7.4). Filters were dried and scintillation cocktail (Meltilex, PerkinElmer) was melted onto the dried filters and allowed to cool to room temperature. Then, filters were placed into cassettes and radioactivity was measured in a Microbeta counter (Molecular Devices). Data were analyzed using “One Site-Total and nonspecific binding” equation using Prism 9. Specific binding was calculated after subtracting the non-specific binding.

### 2.6 Data Analysis

All concentration-response curves were fit to a three-parameter logistic equation in Prism (Graphpad Software, San Diego, CA). BRET2 concentration-response curves were analyzed as either raw net BRET2 (fit Emax-fit Baseline) or by normalizing to a reference agonist for each experiment. Efficacy (Emax) calculations were performed according to Kenakin and colleagues^38^: stimulus-response amplitudes (net BRET2) were normalized to the maximal responding agonist (maximal system response). EC50 and Emax values were estimated from the simultaneous fitting of all biological replicates. Transduction coefficients were calculated as Log(Emax / EC50) as described in Kenakin and colleagues^38^ and propagation of error was conducted at all steps. EC50, Emax, and transduction coefficient values were analyzed first by ANOVAs (F-test of curve fit, one-way ANOVA, or two-way ANOVA as described in the text). *Post hoc* pair-wise comparisons used Tukey-adjusted *p* values to control for multiple comparisons. Significance threshold was set at α = 0.05.

## 3.0 Results

### 3.1 Gα_q_ protein mediated signaling activity potencies for therapeutically promising psychedelics differ between wild-type and polymorphic 5-HT2A receptors

To determine whether naturally occurring, non-synonymous SNPs in the 5-HT_2A_ receptor are likely to affect psychedelic actions *in vivo*, we measured the *in vitro* potencies and efficacies of the wild-type, S12N, T25N, D48N, I197V, A230T, A447V, and H452Y variant 5-HT_2A_ receptors for four therapeutically promising psychedelics. As shown in Figure 2 and Table 1, certain polymorphic 5-HT_2A_ receptors exhibited statistically significant, albeit modest, changes in response to specific psychedelics. The Ala230Thr and His452Tyr receptors displayed a sevenfold decrease in potency for psilocin compared to wild type (38nM vs 5nM, *P*<0.05). The Ala447Val 5-HT_2A_ receptor displayed a threefold increase in potency for 5-MeO-DMT compared to the wild type (3.3nM vs 10.6nM, *P*<0.05). Four polymorphic 5-HT_2A_ receptors exhibited statistically significant changes in mescaline potency. Of these, the largest effect was observed for the Ser12Asn 5-HT_2A_ receptor, which displayed a tenfold increase in potency for mescaline compared to the wild type (340.4 nM vs 3863 nM, *P*<0.0005). None of the SNP-induced alterations in LSD potency were significantly different from the wild-type value (*P*<0.05).

**Figure 2:**
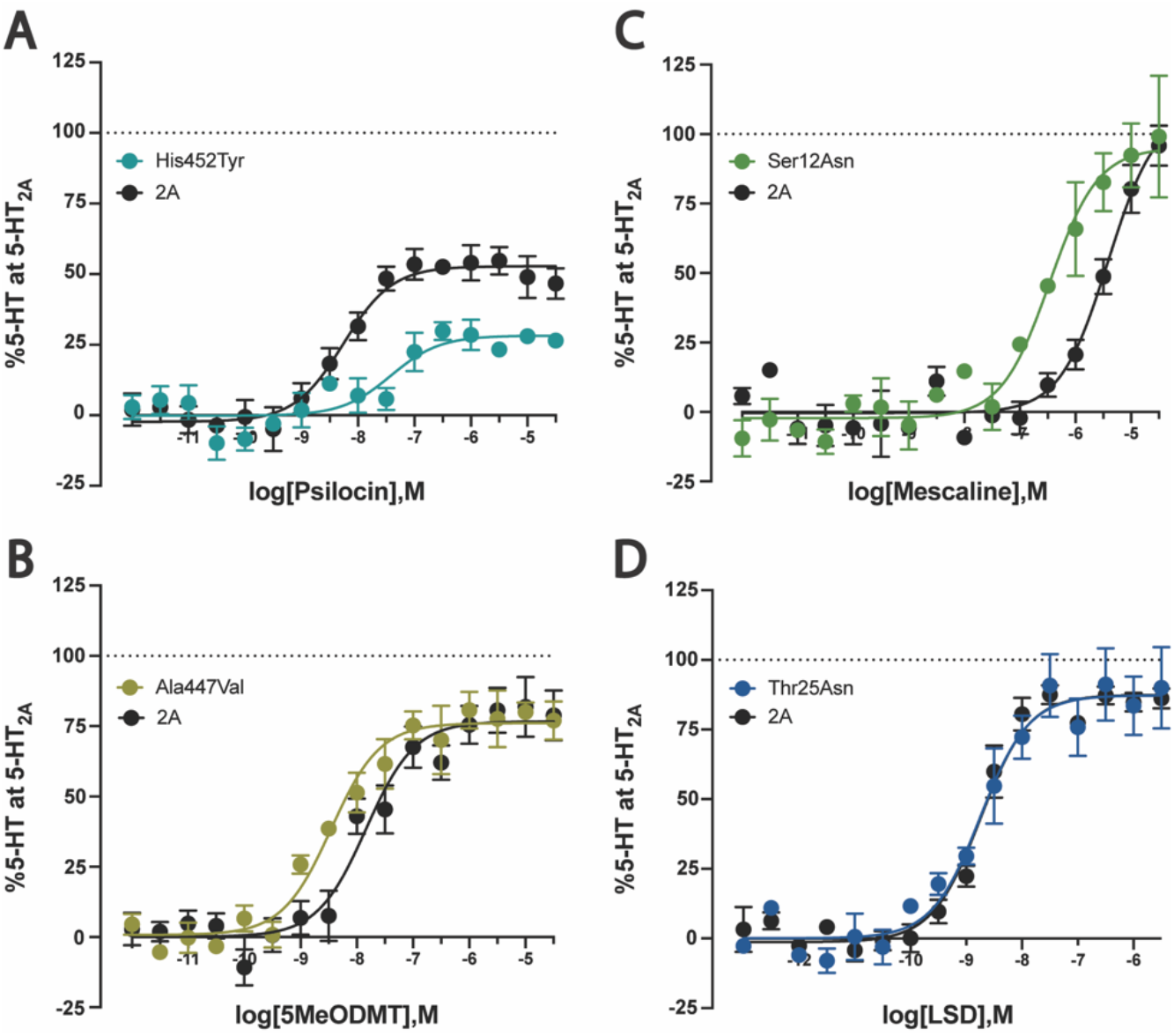
Representative Gq dis concentration-response curves for psychedelic drugs (**a**) Psilocin, (**b**) LSD, (**c**) Mescaline, and (**d**) 5-MeO-DMT at wild-type and polymorphic 5-HT_2A_ receptors using TRUPATH BRET assays. See Table 1 for the complete data matrix.

**Table 1:**
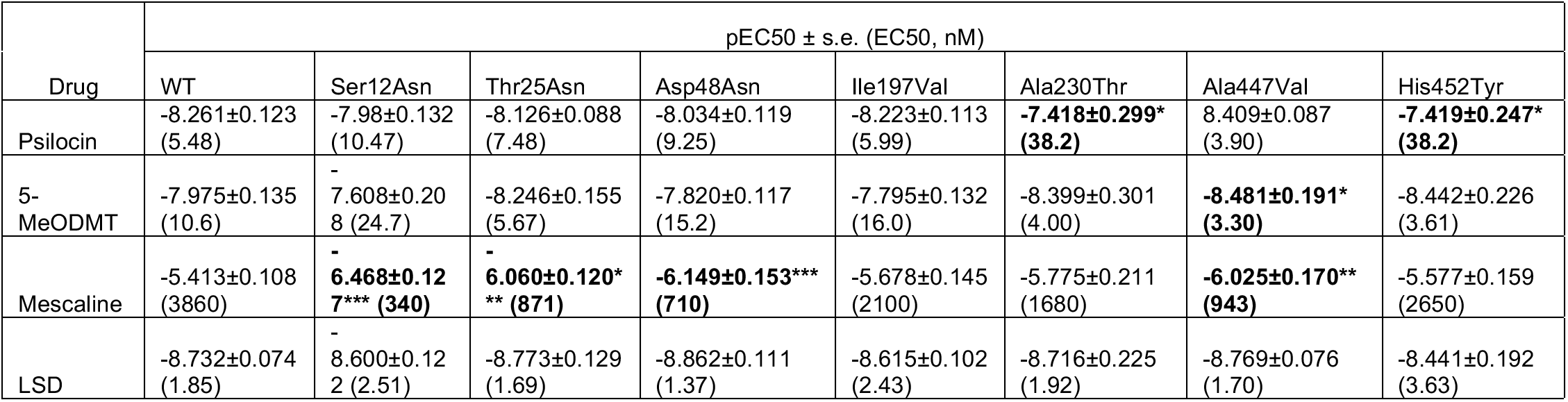
Potency (pEC50) values for several potentially therapeutic psychedelics at wild-type (WT) and polymorphic 5-HT_2A_ receptors as measured using TRUPATH Gq BRET assays. *P<0.05, **P<0.005, ***P<0.0005 by F-test.

### 3.2 βArrestin mediated signaling activity affinities for therapeutically promising psychedelics differ between wild-type and polymorphic 5-HT2A receptors

Having addressed whether several therapeutically promising psychedelics exhibit differential *in vitro* Gα_q_ protein signaling pharmacologies at wild-type and polymorphic 5-HT_2A_ receptors, we next investigated effects on βArrestin recruitment. As shown in Figure 3 and Table 2, the βArr1 potencies of all psychedelics tested were not significantly altered by any of the SNP 5-HT_2A_ receptors tested. In contrast, the βArr2 potencies of 5-MeO-DMT, mescaline, and LSD tested at the Ala230Thr 5-HT_2A_ receptor were significantly altered. Specifically, the 5-MeO-DMT potency was increased twofold compared to wild type, mescaline potency was increased twofold compared to wild type, and LSD potency was increased threefold compared to wild type. As shown in Figure 3 and Table 3, the Ala230Thr 5-HT_2A_ receptor was the only SNP found to cause significant changes in βArrestin2 potencies of 5MeODMT, LSD, and mescaline.

**Figure 3:**
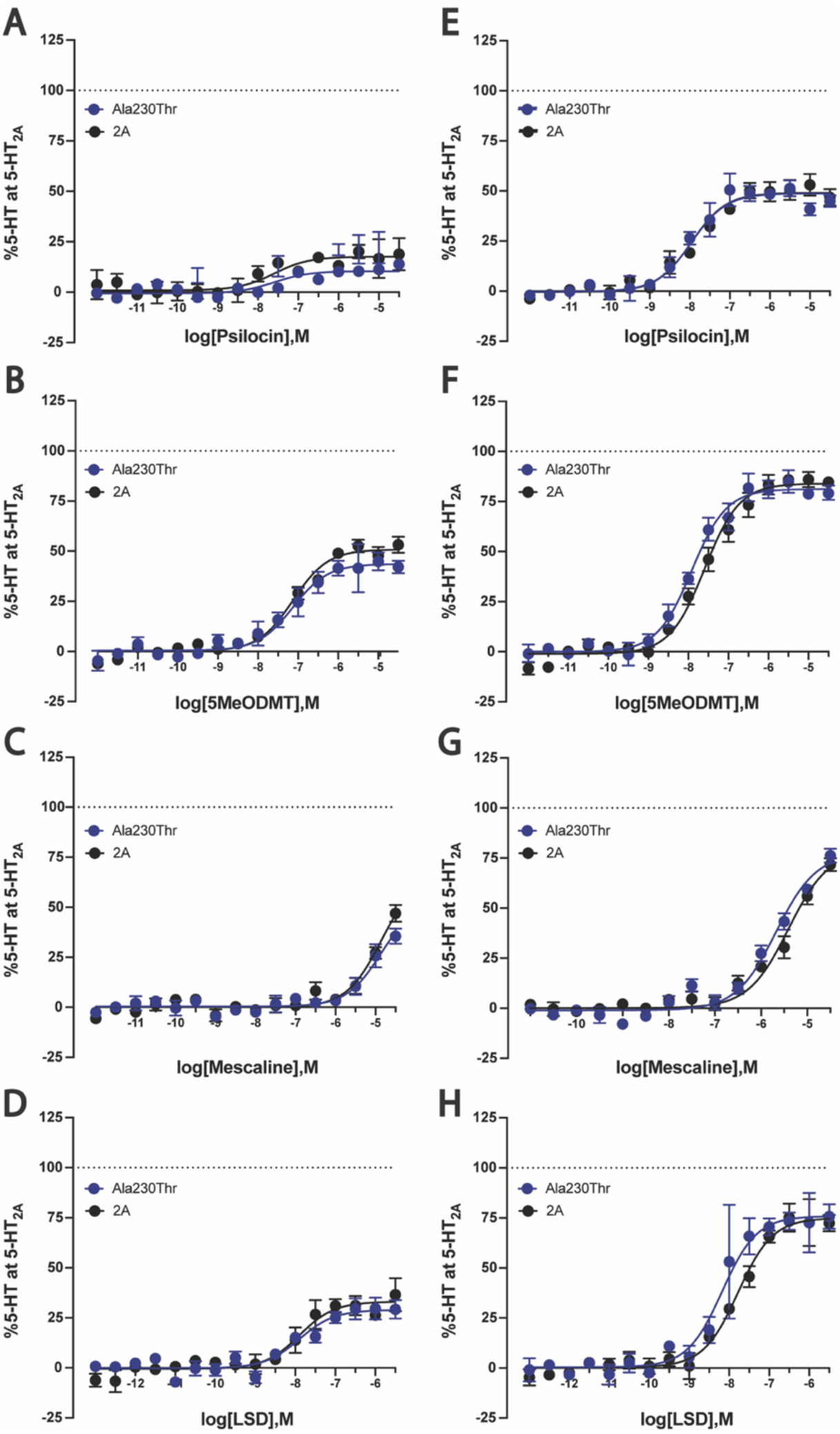
Concentration-response curves for psychedelic drugs (**a**) Psilocin, (**b**) LSD, (**c**) Mescaline, and (**d**) 5-MeODMT at wild-type and polymorphic Ala230Thr 5-HT_2A_ receptors using BRET assays of barrestin-1 and −2 recruitment. See Tables 2 and 3 for the complete data matrix.

**Table 2:** Potency (pEC50) values for several potentially therapeutic psychedelics at wild-type (WT) and polymorphic 5-HT_2A_ receptors as measured using βArr1 recruitment BRET assays.

| Drug | pEC50 ± s.e. (EC50, nM) | | | | | | | |
| --- | --- | --- | --- | --- | --- | --- | --- | --- |
|  | WT | Ser12Asn | Thr25Asn | Asp48Asn | Ile197Val | Ala230Thr | Ala447Val | His452Tyr |
| Psilocin | -7.853±0.352<br>(14.04) | -6.907±0.351<br>(123.9) | -7.631±0.375<br>(23.38) | -7.681±0.302<br>(20.84) | -8.090±0.302<br>(8.13) | -7.101±0.390<br>(79.21) | -7.385±0.256<br>(41.20) | -7.726±0.328<br>(18.81) |
| 5-MeODMT | -7.088±0.073<br>(81.68) | -7.169±0.190<br>(67.75) | -7.073±0.177<br>(84.57) | -7.154±0.123<br>(70.22) | -7.078±0.170<br>(83.48) | -7.222±0.144<br>(59.95) | -7.363±0.325<br>(43.32) | -7.105±0.142<br>(78.44) |
| Mescaline | -4.786±0.162<br>(16365) | -5.421±0.350<br>(3797) | -4.226±0.576<br>(59436) | -5.296±0.211<br>(5057) | -5.062±0.304<br>(8669) | -5.023±0.167<br>(9494) | -4.926±0.334<br>(11869) | -5.408±0.243<br>(3905) |
| LSD | -7.942±0.175<br>(11.42) | -7.742±0.214<br>(18.10) | -7.643±0.186<br>(22.74) | -7.813±0.172<br>(15.37) | -7.780±0.182<br>(16.61) | -7.793±0.164<br>(16.12) | -8.050±0.255<br>(8.92) | -7.583±0.243<br>(26.12) |

**Table 3:** Potency (pEC50) values for several potentially therapeutic psychedelics at wild-type (WT) and polymorphic 5-HT_2A_ receptors as measured using βArr2 recruitment BRET assays. *P<0.05, **P<0.005 by F-test.

| Drug | pEC50 ± s.e. (EC50, nM) | | | | | | | |
| --- | --- | --- | --- | --- | --- | --- | --- | --- |
|  | WT | Ser12Asn | Thr25Asn | Asp48Asn | Ile197Val | Ala230Thr | Ala447Val | His452Tyr |
| Psilocin | -7.812±0.094<br>(15.42) | -8.067±0.155<br>(8.57) | -7.802±0.120<br>(15.77) | -7.611±0.129<br>(24.50) | -7.878±0.094<br>(13.24) | -8.087±0.116<br>(8.18) | -7.810±0.121<br>(15.48) | -7.927±0.097<br>(11.84) |
| 5-MeODMT | -7.595±0.064<br>(25.40) | -7.629±0.094<br>(23.51) | -7.624±0.074<br>(23.75) | -7.647±0.086<br>(22.56) | -7.707±0.088<br>(19.62) | **-7.916±0.071\*\***<br>**(12.13)** | -7.482±0.127<br>(32.98) | -7.523±0.77<br>(30.00) |
| Mescaline | -5.410±0.089<br>(3887) | -5.266±0.118<br>(5421) | -5.383±0.105<br>(4143) | -5.343±0.096<br>(4537) | -5.363±0.112<br>(4333) | **-5.675±0.076\***<br>**(2114)** | -5.258±0.154<br>(5527) | -5.324±0.076<br>(4748) |
| LSD | -7.781±0.086<br>(16.57) | -7.887±0.090<br>(12.97) | -7.924±0.063<br>(11.92) | -7.868±0.88<br>(13.56) | -7.720±0.062<br>(19.05) | **-8.179±0.120\***<br>**(6.62)** | -7.755±0.150<br>(17.59) | -7.641±0.091<br>(22.84) |

### 3.4 Bias calculations for wild-type and polymorphic 5-HT_2A_ receptors

Having addressed whether several SNPs exhibit different *in vitro* signaling for several potentially therapeutic psychedelics, we next quantified changes in bias using the approach of Kenakin and colleagues^38^. To compare between the wild-type and mutant receptors, ΔΔlog(Emax/EC50) values were calculated. As shown in Figure 4, overall biases [ΔΔlog(Emax/EC50) values] for the investigated pathways were significantly altered by multiple SNPs. Psilocin signaling was significantly altered by the Ser12Asn, Ala447Val, and His452Tyr SNPs. The His452Tyr 5-HT_2A_ receptor showed the greatest change in signaling, favoring Gα_q_ signaling over βarr1 and βarr2. The His452Tyr 5-HT_2A_ receptor also demonstrated the largest change in 5-MeO-DMT signaling, increasing Gα_q_ signaling relative to βarr1 and βarr2. Mescaline signaling was significantly altered by the Ser12Asn, Thr25Asn, Asp48Asn, and Ala447Val 5-HT_2A_ receptors. Only the Ala230Thr and His452Tyr 5-HT_2A_ receptors demonstrated altered LSD signaling profiles, favoring Gα_q_ over βarr2 in both cases.

**Figure 4:**
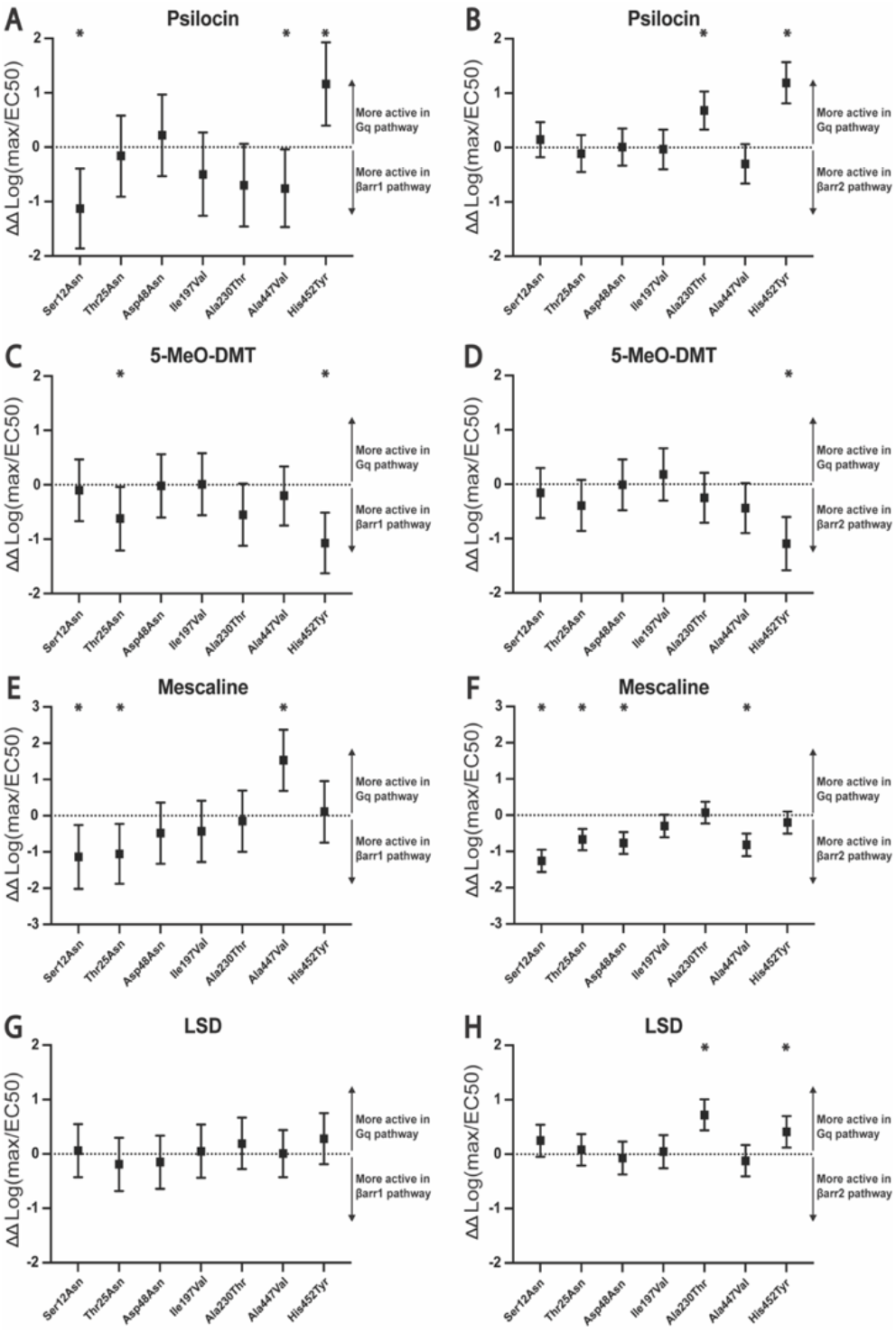
Bias plots showing ΔΔlog(Emax/EC50) values as a measure of ligand bias between signaling pathways. The error bars represent 95% confidence interval; when interval includes zero, the ligands are not biased with respect to each other. (A, B) Psilocin, (C, D) LSD, (E, F) 5-MeO-DMT, (G, H) Mescaline. (A, C, E, G) Gq versus βarr1, (B, D, F, H) Gq versus βarr2.

### 3.5 Effects of 5-HT_2A_ receptor SNPs on receptor expression

To determine whether the 5-HT_2A_ receptor SNPs affected receptor expression, we measured the maximum binding (B_max_) for each by radioligand saturation binding assay as shown in Figure 5 and Table 4. None of the SNPs altered receptor expression more than threefold compared to the wild type. The largest alteration of receptor expression was seen in the Ser12Asn 5-HT_2A_ receptor which was, on average, reduced less than threefold relative to the wild-type receptor. As we previously demonstrated our BRET-based assays are relatively insensitive to differences in receptor expression^39^. We note that even though the Ser12Asn and Ala230Thr polymorphisms were both expressed at levels higher than WT no uniform differences in agonist potencies or efficacies were observed. Thus, the differences observed between the *in vitro* pharmacology of the seven variant 5-HT_2A_ receptors tested probably do not arise due to differential expression.

**Figure 5:**
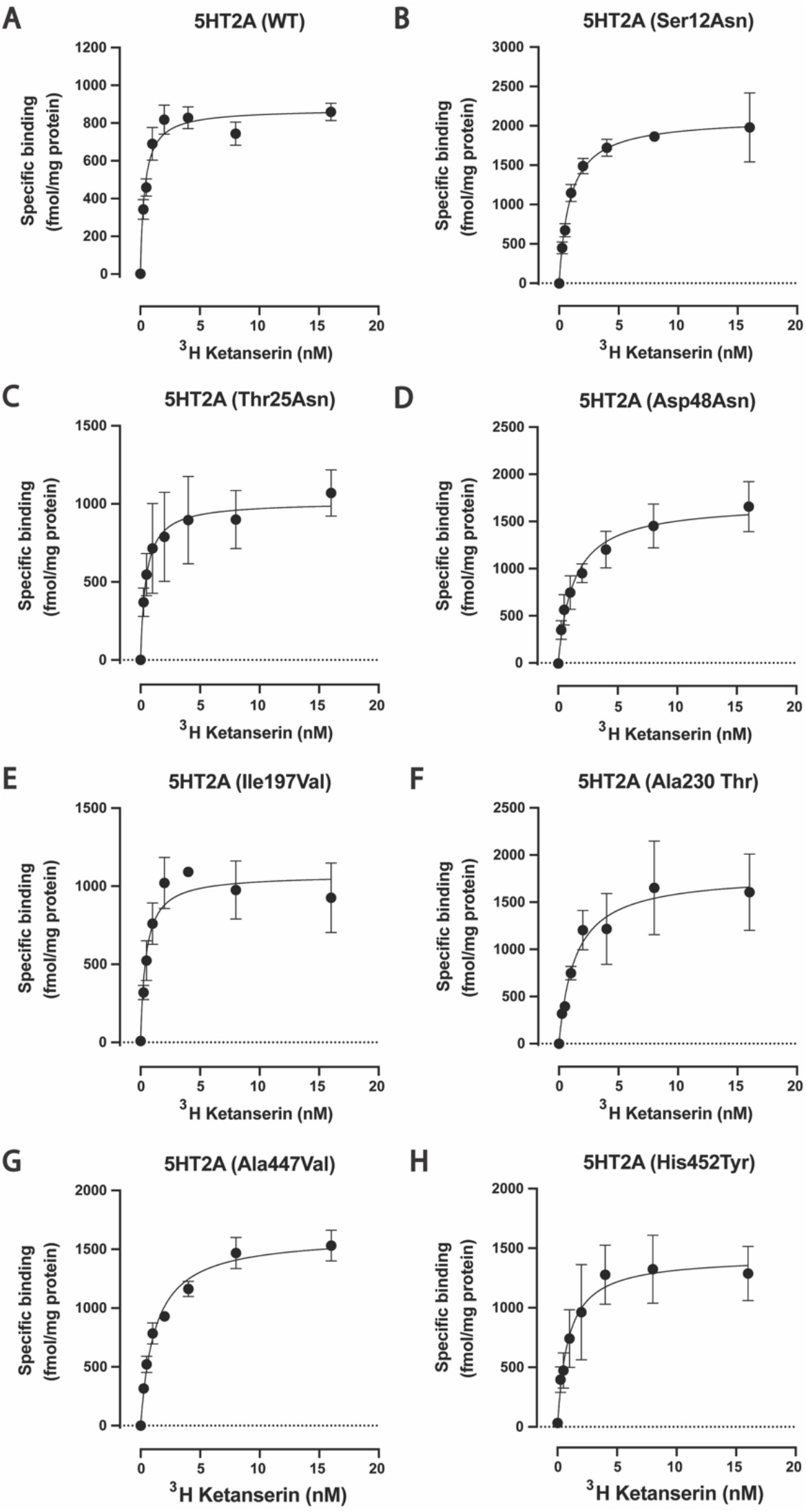
Saturation radioligand binding analysis of WT and polymorphic 5HT2A receptors as a measure of receptor expression. (A) WT 5HT2A, (B) Ser12Asn, (C) Thr25Asn, (D) Asp48Asn, (E) Ile197Val, (F) Ala230Thr, (G) Ala447Val, (H) His452Tyr.

**Table 4:** Bmax values at wild type (WT) and polymorphic 5-HT_2A_ receptors as measured using radioligand competition binding

|  | WT | Ser12Asn | Thr25Asn | Asp48Asn | Ile197Val | Ala230Thr | Ala447Val | His452Tyr |
| --- | --- | --- | --- | --- | --- | --- | --- | --- |
| Bmax (fmol/mg) | 874.4±38.4 | 2100±113 | 1013±120 | 1698±148 | 1073±91.1 | 1802±215 | 1618±72.4 | 1429±185 |

## 4.0 Discussion

The major finding of this study is that certain 5-HT_2A_ receptor non-synonymous SNPs can alter the *in vitro* pharmacology of the tested psychedelics. As we have demonstrated in the current study, even polymorphisms some distance from the orthosteric binding site of the receptor can have substantial effects on the *in vitro* pharmacology of a given drug. Given that several psychedelic drugs are in clinical trials, this information could aid in the design and ultimate interpretation of clinical studies.

Our prior study^36^ examined some of these polymorphisms on the agonist and antagonist actions of several drugs, although functional selectivity was not. Additionally, of the compounds tested in the current study, only 5-MeO-DMT was examined. Based on the current results, our data imply that individuals carrying the Ala230Thr variant might show altered responses to all of the psychedelics studied given that Gα_q_ and/or βarr2 signaling was altered. Additionally, we would predict that individuals carrying the His452Tyr variants might show altered responses to psilocin and psilocybin, as large decreases in psilocin Gα_q_ potency were found to significantly alter the signaling bias. For mescaline, the most dramatic change was seen at the Ser12Asn polymorphism with a tenfold increase in potency. Of note, our study only examined binding potency and efficacy at a single five-minute timepoint. βarr2 signaling of LSD, for instance, has been shown to increase in potency and efficacy over time^18^. Therefore, future studies could examine the effects of 5-HT_2A_ receptor SNPs on signaling kinetics.

In summary, our findings indicate that 5-HT_2A_ receptor SNPs can alter the *in vitro* pharmacology of some therapeutically promising psychedelics. Prior studies have already demonstrated that the His452Tyr as well as Thr25Asn and Ile197Val 5-HT_2A_ receptor polymorphisms can exhibit pronounced differences from the wild-type receptor in the pharmacologies of atypical antipsychotics^36^. Our results confirm that these receptors can exhibit pronounced differences from the wild-type receptor in the pharmacologies of psychedelics. Additionally, we find that the polymorphisms Ser12Asn, Ala230Thr, and Ala447Val 5-HT_2A_ receptor polymorphisms also exhibit pronounced differences. These results may have relevance for the design and interpretation of future clinical trials.

## ACKNOWLEDGEMENTS

This work was supported by R37DA04567 from the National Institute of Drug Abuse to BLR. GS was supported by the Medical Scientist Training Program at UNC.

## Notes

### Competing Interest Statement

The authors have declared no competing interest.

